# Breathing interoception of the anterior cingulate cortex in common marmosets

**DOI:** 10.64898/2026.11.23.768718

**Authors:** Shahriar SheikhBahaei, Gurueswar Nagarajan, Mitchell Bishop, Yogita Chudasama

**Affiliations:** Department of Neurobiology & Behavior, Stony Brook University Renaissance School of Medicine, Stony Brook, NY 11794, USA; Center for Nervous System Disorders, Stony Brook University, Stony Brook, NY 11794, USA; Section on Behavioral Neuroscience, National Institute of Mental Health, Bethesda, MD 20892, USA; Department of Neurobiology, University of Pittsburgh School of Medicine, Pittsburgh, PA, United States

**Keywords:** anterior cingulate cortex, breathing, common marmoset, interoception, non-human primates

## Abstract

Breathing links brainstem homeostatic circuits with cortical interoceptive processing: its fundamental rhythm is generated in the brainstem, continuously monitored by the suprabrainstem regions, and rapidly modulated by emotionally salient states [1–10]. Although the anterior cingulate cortex (ACC) has been implicated in autonomic control and the affective representation of bodily states, its causal role in regulating respiratory states remains unknown [11–14]. Here, we show that early-life disruption of the ACC in common marmosets selectively blunts the ventilatory response to a fear-evoking stimulus later in development, while preserving the overall pattern of breathing dynamics. These findings demonstrate that the primate ACC is required to adapt breathing to emotionally salient situations.

## Main text

Interoception comprises the processes by which the brain monitors, interprets, and regulates the body’s physiological state [1,2,4,5,15]. Breathing is a particularly informative interoceptive signal, bridging homeostatic regulation and emotional processing [7–9,16–21]. Respiratory signals are conveyed through partially separable discriminative and affective pathways [9,19]. Attention to breathing engages a distributed cortical network that includes the anterior cingulate, insular, premotor, and hippocampal regions [7]. Conversely, emotional and cognitive states can likewise dynamically modulate spontaneous breathing through limbic, paralimbic, and brainstem circuits [8]. Together, these observations place the cingulate cortex in a broader forebrain network that integrates respiratory signals with emotional context to shape perception and behavior [16,22–25].

Within this broader respiratory-interoceptive network, the anterior cingulate cortex (ACC) links bodily states to emotional significance. Human imaging and lesion studies implicate ACC regions in autonomic regulation and allostatic control [11–13], and in nonhuman primates, perturbing the ACC alters sustained arousal and threat-related cardiovascular and behavioral responses [14,26]. Together, these findings suggest that ACC circuits coordinate physiological responses to emotionally salient events.

However, whether the ACC broadly regulates breathing or selectively couples emotional salience to respiratory control is unknown. The common marmoset provides a valuable nonhuman primate model to address this question because the anatomical organization of the ACC has been well characterized in this species [27]. Furthermore, early-life disruption of this region impairs social vocal maturation, a process heavily dependent on breathing coordination [28], and alters the structure of downstream vocal networks [29]. We therefore asked whether early-life ACC disruption also alters the respiratory response to fear-relevant stimuli later in development.

To test this hypothesis, infant marmosets underwent early ACC disruption (n = 4) or sham surgery (n = 3) at postnatal days 14-16 [29] and were studied months later during adolescence (mean age: 11 months; **Figure 1A**). Structural magnetic resonance imaging and histological analysis confirmed substantial disruption of area 24a and 24b, including reduced anterior cingulate volume and loss of the normal radial organization of myelinated fibers [29] **(Figure 1B**). Before assessing respiratory responses, we first established that ACC-disrupted animals retained a defensive response to the fear-relevant stimulus. Animals were presented with a neutral object (e.g., toy car) and a raptor model (a fake owl) previously shown to evoke antipredator behavior in marmosets [30]. Defensive reactions were quantified from video recordings by an investigator blinded to the animals’ group assignment. The owl elicited a robust threat response marked by several defensive indicators and heightened vigilance (**Figure 1C)**, significantly increasing the composite threat-response score across animals compared with the neutral object (p = 0.017; **Figure 1D**). Importantly, threat reactivity did not differ between ACC-disrupted and control animals during either neutral-object (p = 0.8) or owl presentation (p = 0.6). Thus, early ACC disruption did not abolish behavioral sensitivity to threat.

**Figure 1.**
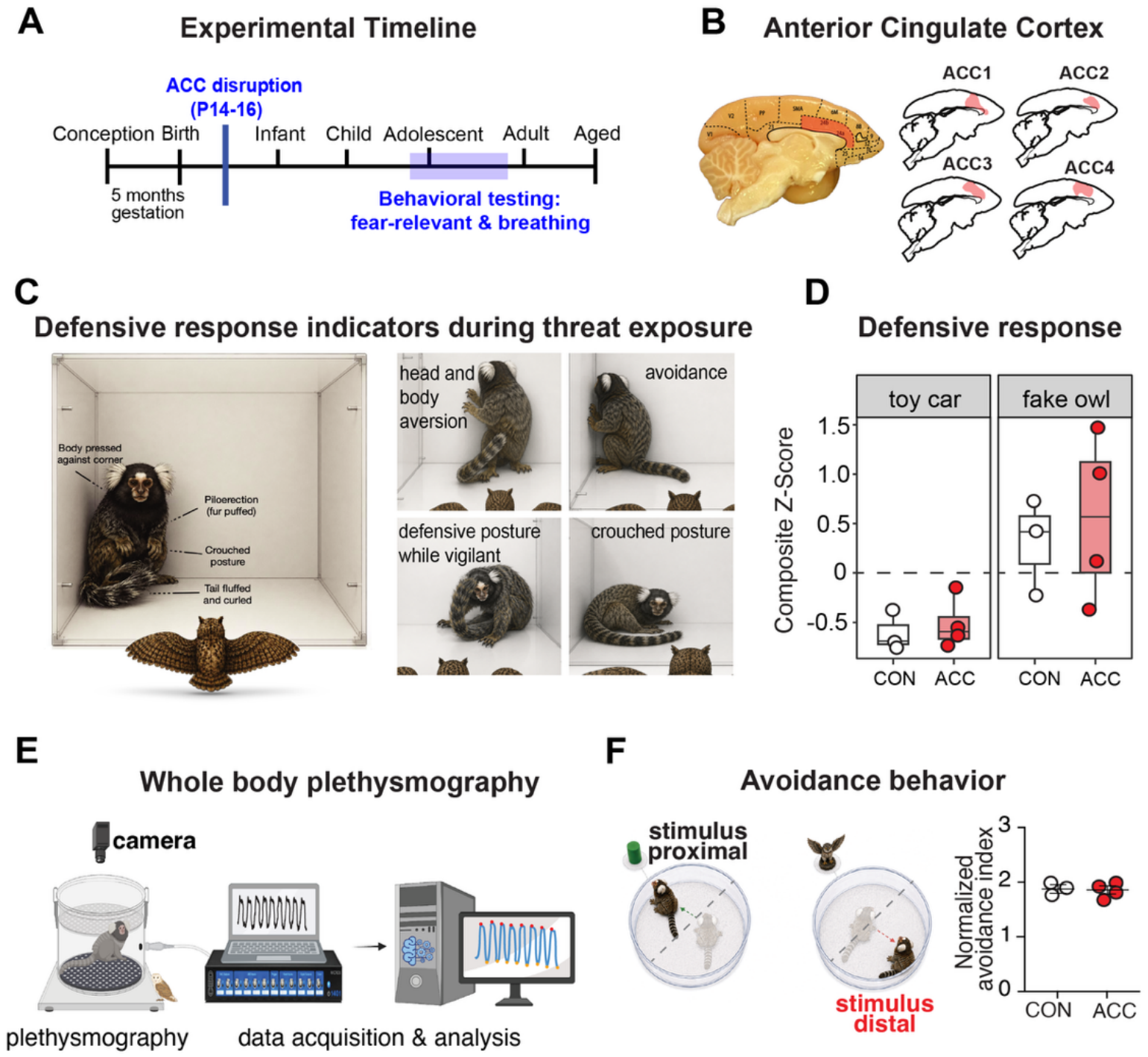
Early anterior cingulate cortex disruption elicits defensive reactions to a fear-relevant stimulus. **(A)** Longitudinal experimental timeline. Common marmosets underwent early anterior cingulate cortex (ACC) disruption or sham control surgery at postnatal days 14–16 and were subsequently tested during adolescence using behavioral assessment and awake whole-body plethysmography. **(B)** Location of anterior cingulate cortex (ACC) and extent of ACC disruption in individual animals (ACC1-ACC4). Anatomical divisions and nomenclature [37] superimposed onto sagittal view of marmoset brain. Shaded region indicates affected ACC territory (24a, 24b). Detailed histological verification and characterization of the extent of ACC disruption in these animals have been reported previously [28]. Abbreviations: V1, visual area 1; V2, visual area 2; PP, posterior parietal; SMA, supplementary motor area; 6M, medial area 6 of cortex; 8B/9/10/32/25 & 14 areas of cortex. **(C)** Schematic of behavioral assessment and defensive indicators used to quantify threat-like responses to the fear-evoking owl stimulus, including defensive posture, piloerection, crouching, avoidance, and vigilance (focused looking at the stimulus). Schematics were generated with assistance of ChatGPT-6, AI generative tools within BioRender, and Adobe Illustrator 2027. **(D)** Overall threat response towards a neutral object (toy car) and a fear-evoking object (fake owl). The composite Z-scores integrate three standardized behavioral measures: severity-weighted duration of defensive behaviors, frequency of defensive bouts, and duration of vigilance toward the stimulus. Each measure was independently converted to a Z-score, and the three Z-scores were averaged to generate a composite score for each animal. A composite Z-score greater than 0 indicates increased defensive reactivity. Data are presented as box plots displaying the median and interquartile range for Controls (CON, white) and ACC (red) groups. The horizontal dashed black line represents the overall mean (Z = 0). The presence of the fake owl significantly elevated the composite defensive score across all subjects (p = 0.017), with no group differences in reactivity toward the neutral toy car (p = 0.8) or the threatening fake owl (p = 0.6). Points represent individual animals. Statistical comparisons were performed using a linear mixed-effects model followed by pairwise comparisons with the Kenward–Roger approximation. Together, these measures indicate that early life ACC disruption did not abolish the behavioral response to the fear-evoking stimulus. **(E)** Schematic of the awake whole-body plethysmography setup used for respiratory recording, data acquisition, and automated extraction of respiratory features. Schematics were generated using BioRender.com. **(F)** Schematic of avoidance behavior during presentation of the fear-evoking owl within the plethysmography chamber. We quantified an avoidance index from relative time spent in stimulus-proximal and stimulus-distal zones and normalized it to the neutral (toy) object condition (See Methods). Each point represents one animal. Schematics were generated using BioRender.com. Horizontal bars indicate mean ± SEM.

We next examined whether this preserved behavioral response was accompanied by breathing modulation. Breathing was recorded using awake whole-body plethysmography (**Figure 1E**). Following a 40-min baseline recording, animals were exposed, sequentially, to a neutral object, a fear-evoking owl, and a 10-min hypoxic challenge. This design enabled a within-subject comparison of respiratory responses to affectively neutral and fear-relevant stimuli while using hypoxia as a homeostatic respiratory challenge. Object presentations lasted 30 s and were separated by a 60-s interstimulus interval. Breathing was quantified using an automated workflow for plethysmographic data acquisition and respiratory feature extraction [31].

To confirm that the owl stimulus remained behaviorally salient within the plethysmography chamber, we quantified avoidance behavior during presentation of the neutral object and the owl object. Behavioral responses were scored from video recordings by an investigator blinded to the group. An avoidance index (AI) was calculated from the relative time spent in the stimulus-proximal and stimulus-distal zones of the chamber (**Figure 1F**), and owl-evoked avoidance was expressed as a change relative to the neutral object condition (ΔAI=AI_owl_™ AI_neutral_). Positive ΔAI values indicate greater avoidance of the owl than the neutral object. Both sham and ACC-disrupted animals showed increased avoidance of the owl relative to the neutral object (**Figure 1F**), confirming that the owl remains behaviorally threatening under the same experimental condition used to measure breathing.

We then asked whether early ACC disruption altered breathing under low-salience conditions. During presentation of the neutral object, respiratory trajectories remained close to baseline in both groups. Animals with early ACC disruption showed changes in mean breathing rate (respiratory frequency), tidal volume (breath depth), and minute ventilation (air movement in lungs) that were indistinguishable from sham controls (**Figures 2A-C**, white panels). Therefore, early ACC disruption neither produced an overt baseline ventilatory abnormality nor an altered respiratory response to a behaviorally neutral stimulus.

**Figure 2.**
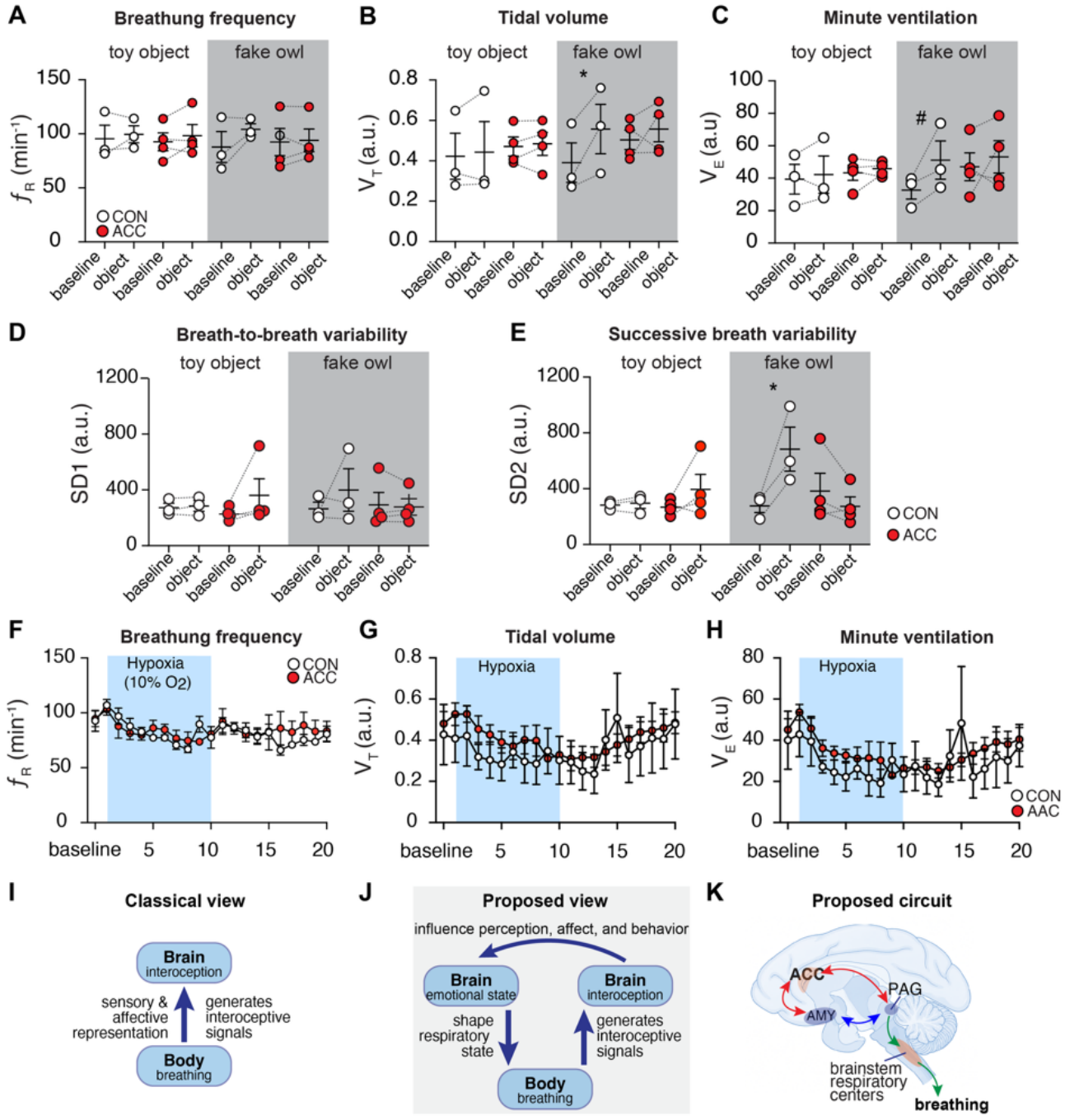
Early ACC disruption selectively attenuates respiratory responses to fear-evoking stimulus in marmosets, while preserving homeostatic responses to hypoxia. (**A-C**) Respiratory responses to presentation of a neutral toy object and a fear-evoking fake owl in control (CON, white) and ACC-disrupted (ACC, red) animals. Respiratory frequency (*f*_R_), tidal volume (*V*_T_), and minute ventilation (*V*_E_) are shown during the baseline period immediately preceding each stimulus and during object presentation. The fake owl increased tidal volume and minute ventilation in control animals, whereas these responses were attenuated following early ACC disruption. Data are shown as mean ± SEM. * *P* = 0.046; #*P* = 0.07. *p*-values denote group comparisons of object-evoked respiratory change; paired *t*-test. (**D-E**) Breathing variability during toy object and owl-object trials, quantified by Poincaré-derived SD1 and SD2. SD1 reflects short-timescale breath-to-breath variability, whereas SD2 reflects slower variability in respiratory timing across successive breaths. Exposure to fake owl increased respiratory variability in controls, particularly SD2, an effect that was attenuated following ACC disruption. * *P* = 0.057 (**F-H**) Respiratory responses to a 10-min hypoxic challenge (10% O_2_, blue shading). Respiratory frequency, tidal volume, and minute ventilation are shown before (baseline), during, and after hypoxia. ACC-disrupted and control animals showed broadly similar respiratory trajectories during the hypoxic challenge and subsequent recovery, confirming that early-life ACC disruption did not produce a generalized impairment in homeostatic respiratory control. (**I-K**) Proposed role of ACC in emotion-dependent regulation of breathing. **(I)** Classical models of respiratory interoception emphasize ascending signals from the body to the brain, whereby breathing generates interoceptive signals that contribute to sensory and affective representations of bodily state. **(J)** Our findings support a complementary bidirectional framework in which forebrain emotional circuits not only receive and interpret respiratory signals but also shape respiratory state according to emotional context. The resulting changes in breathing, in turn, generate interoceptive signals that can influence perception, affect, and behavior. **(K)** Proposed circuitry through which the ACC may contribute to emotion-dependent respiratory regulation. Interactions among the ACC, amygdala (AMY), and periaqueductal gray (PAG) provide potential descending routes through which emotional salience can modulate brainstem respiratory centers and breathing. This model positions the ACC within a broader forebrain–brainstem circuit that couples emotional state to respiratory output while preserving brainstem mechanisms supporting homeostatic respiratory control.

A different pattern emerged when the stimulus carried affective salience. During presentation of the fear-evoking owl, sham controls exhibited a robust ventilatory response driven primarily by a 44% increase in tidal volume (0.39 to 0.56 a.u., *P* = 0.046, paired *t*-test). Mean breathing rate also increased from 87 to 103 min^-1^ (*P* = 0.21, paired *t*-test), and minute ventilation rose from 33 to 51 a.u. (*P* = 0.072, paired *t*-test; **Figures 2A-C**, gray panels). In contrast, ACC-disrupted animals did not show the same coordinated ventilatory response. During fear-object exposure, these animals showed only a small increase in tidal volume (0.50 to 0.56 a.u.; *P* = 0.5, paired *t*-test) and essentially no increase in respiratory frequency (92 to 94 min^-1^; *P* = 0.76, paired *t*-test), so their increase in minute ventilation was much smaller (47 to 53 a.u.; *P* = 0.2, paired *t*-test, **Figures 2B-C**). Thus, although ACC-disrupted animals retained the capacity to generate breaths of similar magnitude to sham controls, their fear-linked ventilatory response was markedly attenuated. This breathing pattern indicates that early-life ACC disruption selectively impairs the normal recruitment of ventilatory responses by emotional salience, rather than the capacity to generate increased ventilation *per se*.

We next asked whether ACC disruption altered variability of breathing. To address this, we quantified respiratory variability using Poincaré analysis and observed the same overall pattern. During the neutral-object presentation, short-timescale variability (SD1) was broadly comparable between ACC-disrupted and sham controls (**Figure 2D**). During fear-object exposure, however, SD1 increased in sham animals (265 to 399 a.u.) but showed little overall change following early-life ACC disruption (from 292 to 277 a.u.). The group difference was even more pronounced in SD2, which captures longer-timescale respiratory dynamics. During neutral-object presentation, SD2 changed little in the controls but increased modestly in the ACC group. During fear-object exposure, however, controls showed a marked increase in SD2 (273 to 677 a.u.) whereas ACC-disrupted animals instead showed a decrease (383 to 271 a.u.; ΔSD2 _fear-object_ = ™110 in ACCx vs. +407 in sham; *P* = 0.057, Mann–Whitney U test; **Figure 2E)**. These data indicate that early-life ACC disruption attenuates not only the ventilatory response to emotionally salient stimuli but also the longer timescale organization of breathing associated with threat. In other words, the deficit reflects more than reduced breath depth; it represents a broader disruption of emotionally dependent respiratory dynamics.

We then asked whether the attenuated fear response reflected a selective loss of emotion-dependent respiratory modulation or a broader impairment of respiratory control. We therefore challenged the same animals with 10% oxygen, a homeostatic stimulus that engages respiratory control independently of emotional context. Animals with early ACC disruption and sham controls showed broadly similar responses across respiratory frequency, tidal volume, and minute ventilation. Consistent with the characteristic hypoxic-evoked breathing previously described in awake marmosets [31], these same respiratory markers declined during hypoxia (**Figures 2F-H**, blue panels). They recovered toward baseline following the return to normoxia in both groups (**Figures 2F-H**, white panels). Thus, early-life ACC disruption did not produce a generalized deficit in homeostatic respiratory control, supporting the conclusion that the neonatal disruption of the ACC selectively contributes to emotion-dependent modulation of breathing in later adolescence.

Together, these findings indicate that core features of homeostatic respiratory control remained largely intact after early ACC disruption, whereas the ventilatory response to a fear-evoking stimulus was selectively attenuated. This developmental phenotype, therefore, reflects a specific impairment in emotion-dependent respiratory modulation rather than a generalized deficit in breathing. Our findings identify the ACC as part of a forebrain circuit that links emotional salience to respiratory output, enabling affectively meaningful stimuli to recruit changes in ventilation [6,7,13]. Whereas recent studies have emphasized perceptual, predictive, and anxiety-related aspects of breathing, particularly within insular cortical networks [9,16], our study addresses a complementary question: whether a primate forebrain region is required to transform emotionally salient stimuli into respiratory responses. Our results identify ACC as one such region (**Figure 2J-K)**. Because both sham and ACC-disrupted animals showed comparable avoidance of the owl stimulus, the attenuated ventilatory response following ACC-disruption is unlikely to reflect impaired detection of, or behavioral responsiveness to, the predator-like stimulus.

Prior human studies have largely focused on how breathing-related signals are perceived, predicted, or misinterpreted in intact adults, particularly in the context of anxiety and interoceptive processing [9,16]. Although these works established breathing as a major interoceptive signal, it left unresolved whether a primate cortical region is required to couple emotionally salient stimuli with respiratory responses. Our data address this question by shifting the focus from respiratory perception to respiratory regulation. Early ACC disruption selectively attenuated the ventilatory response to fear while sparing homeostatic respiratory control, identifying the ACC as a component of the forebrain circuitry through which emotional salience shapes breathing.

The preserved behavioral response to the owl helps constrain the interpretation of the respiratory deficit. Animals with early life ACC disruption did not behave as if the stimulus was undetected or lacked emotional significance. Instead, the impairment was selective to the respiratory response that normally accompanies threat. This interpretation aligns with earlier work in the same developmental model, which showed that early-life ACC disruption alters GABA-positive neurons in the basomedial amygdala and dorsal periaqueductal gray [29]. Their established roles in defensive behavior, autonomic regulation, and vocal-respiratory patterning [14,29], suggest that the present findings reflect disrupted maturation of a broader cingulate–amygdala–periaqueductal pathway that links emotional salience to respiratory control.

These findings also broaden how breathing interoception can be conceptualized. Much of the field has emphasized the ascending dimension of interoception: how cortical networks sense, predict, and assign affective significance to respiratory signals [9,16,23–25,32,33]. Our findings highlight the complementary descending dimension: how forebrain circuits shape respiratory state before those signals are available for interoceptive processing. In this view, breathing should be viewed not only as an interoceptive signal that informs the brain, but also as a physiological state actively shaped by forebrain circuits before it is sensed (Figure 3A). Moreover, our results support a recently proposed model in which internal state is described as a bidirectional “infinity loop” between the body and brain [34]. These perspectives may be particularly relevant to developmental neuropsychiatric disorders, in which abnormal arousal, anxiety, or bodily-state regulation could arise from disrupted coupling between affective circuits and respiratory/autonomic output, rather than from primary dysfunction of brainstem respiratory control [35,36].

## Acknowledgments

This research was supported by the Intramural Research Program of the National Institute of Mental Health (ZIA MH002951 to YC), part of the National Institutes of Health (NIH). The contributions of the NIH author(s) are considered Works of the United States Government. The findings and conclusions presented in this paper are those of the author(s) and do not necessarily reflect the views of the NIH or the U.S. Department of Health and Human Services. We thank Dr. Denis Matrov for help with lesion reconstruction. GN is now at The Henry Jackson Foundation for the Advancement of Military Medicine, Inc., Bethesda, MD, USA. MB is currently a graduate student at the University of Pittsburgh.

## References

1. Chen, W.G., Schloesser, D., Arensdorf, A.M., Simmons, J.M., Cui, C., Valentino, R., Gnadt, J.W., Nielsen, L., Hillaire-Clarke, C.S., Spruance, V., et al. (2021). The Emerging Science of Interoception: Sensing, Integrating, Interpreting, and Regulating Signals within the Self. Trends Neurosci. 44, 3–16.

2. Khalsa, S.S., Adolphs, R., Cameron, O.G., Critchley, H.D., Davenport, P.W., Feinstein, J.S., Feusner, J.D., Garfinkel, S.N., Lane, R.D., Mehling, W.E., et al. (2018). Interoception and mental health: A roadmap. Biol. Psychiatry Cogn. Neurosci. Neuroimaging 3, 501–513.

3. Quadt, L., Critchley, H.D., and Garfinkel, S.N. (2018). The neurobiology of interoception in health and disease. Ann. N. Y. Acad. Sci. 1428, 112–128.

4. Critchley, H.D., and Garfinkel, S.N. (2017). Interoception and emotion. Curr. Opin. Psychol. 17, 7–14.

5. Craig, A.D. (2002). How do you feel? Interoception: the sense of the physiological condition of the body. Nat. Rev. Neurosci. 3, 655–666.

6. Davenport, P.W., and Vovk, A. (2009). Cortical and subcortical central neural pathways in respiratory sensations. Respir. Physiol. Neurobiol. 167, 72–86.

7. Herrero, J.L., Khuvis, S., Yeagle, E., Cerf, M., and Mehta, A.D. (2018). Breathing above the brain stem: volitional control and attentional modulation in humans. J. Neurophysiol. 119, 145–159.

8. Evans, K.C., Dougherty, D.D., Schmid, A.M., Scannell, E., McCallister, A., Benson, H., Dusek, J.A., and Lazar, S.W. (2009). Modulation of spontaneous breathing via limbic/paralimbic-bulbar circuitry: an event-related fMRI study. Neuroimage 47, 961–971.

9. Chan, P.-Y.S., Lee, L.-Y., and Davenport, P.W. (2024). Neural mechanisms of respiratory interoception. Auton. Neurosci. 253, 103181.

10. Sheikhbahaei, S., and Smith, J.C. (2017). Breathing to inspire and arouse. Science 355, 1370–1371.

11. Critchley, H.D., Mathias, C.J., Josephs, O., O’Doherty, J., Zanini, S., Dewar, B.-K., Cipolotti, L., Shallice, T., and Dolan, R.J. (2003). Human cingulate cortex and autonomic control: converging neuroimaging and clinical evidence. Brain 126, 2139–2152.

12. Kleckner, I.R., Zhang, J., Touroutoglou, A., Chanes, L., Xia, C., Simmons, W.K., Quigley, K.S., Dickerson, B.C., and Barrett, L.F. (2017). Evidence for a Large-Scale Brain System Supporting Allostasis and Interoception in Humans. Nat. Hum. Behav. 1.

13. Ferraro, S., Klugah-Brown, B., Tench, C.R., Bazinet, V., Bore, M.C., Nigri, A., Demichelis, G., Bruzzone, M.G., Palermo, S., Zhao, W., et al. (2022). The central autonomic system revisited - Convergent evidence for a regulatory role of the insular and midcingulate cortex from neuroimaging meta-analyses. Neurosci. Biobehav. Rev. 142, 104915.

14. Rudebeck, P.H., Putnam, P.T., Daniels, T.E., Yang, T., Mitz, A.R., Rhodes, S.E.V., and Murray, E.A. (2014). A role for primate subgenual cingulate cortex in sustaining autonomic arousal. Proc Natl Acad Sci USA 111, 5391–5396.

15. Greenwood, B.M., and Garfinkel, S.N. (2025). Interoceptive mechanisms and emotional processing. Annu. Rev. Psychol. 76, 59–86.

16. Harrison, O.K., Köchli, L., Marino, S., Luechinger, R., Hennel, F., Brand, K., Hess, A.J., Frässle, S., Iglesias, S., Vinckier, F., et al. (2021). Interoception of breathing and its relationship with anxiety. Neuron 109, 4080–4093.e8.

17. Weng, H.Y., Feldman, J.L., Leggio, L., Napadow, V., Park, J., and Price, C.J. (2021). Interventions and manipulations of interoception. Trends Neurosci. 44, 52–62.

18. Ashhad, S., Kam, K., Del Negro, C.A., and Feldman, J.L. (2022). Breathing rhythm and pattern and their influence on emotion. Annu. Rev. Neurosci. 45, 223–247.

19. Del Negro, C.A., Funk, G.D., and Feldman, J.L. (2018). Breathing matters. Nat. Rev. Neurosci. 19, 351–367.

20. Turk, A.Z., Bishop, M., Adeck, A., and SheikhBahaei, S. (2022). Astrocytic modulation of central pattern generating motor circuits. Glia 70, 1506–1519.

21. Sheikhbahaei, S., Gourine, A.V., and Smith, J.C. (2017). Respiratory rhythm irregularity after carotid body denervation in rats. Respir. Physiol. Neurobiol. 246, 92–97.

22. Chemis, I.M., Köchli, L., Marino, S., Russell, B.R., Stephan, K.E., and Harrison, O.K. (2025). The relationship between interoception of breathing, anxiety, and resting-state functional connectivity in the brain. Cogn. Affect. Behav. Neurosci. 25, 1795–1806.

23. Peiffer, C., Poline, J.B., Thivard, L., Aubier, M., and Samson, Y. (2001). Neural substrates for the perception of acutely induced dyspnea. Am. J. Respir. Crit. Care Med. 163, 951–957.

24. Binks, A.P., Evans, K.C., Reed, J.D., Moosavi, S.H., and Banzett, R.B. (2014). The time-course of cortico-limbic neural responses to air hunger. Respir. Physiol. Neurobiol. 204, 78–85.

25. Stoeckel, M.C., Esser, R.W., Gamer, M., Büchel, C., and von Leupoldt, A. (2016). Brain Responses during the Anticipation of Dyspnea. Neural Plast. 2016, 6434987.

26. Alexander, L., Wood, C.M., Gaskin, P.L.R., Sawiak, S.J., Fryer, T.D., Hong, Y.T., McIver, L., Clarke, H.F., and Roberts, A.C. (2020). Over-activation of primate subgenual cingulate cortex enhances the cardiovascular, behavioral and neural responses to threat. Nat. Commun. 11, 5386.

27. Schaeffer, D.J., Gilbert, K.M., Ghahremani, M., Gati, J.S., Menon, R.S., and Everling, S. (2019). Intrinsic functional clustering of anterior cingulate cortex in the common marmoset. Neuroimage 186, 301–307.

28. Subramanian, H.H., and Holstege, G. (2010). Periaqueductal gray control of breathing. Adv. Exp. Med. Biol. 669, 353–358.

29. Nagarajan, G., Matrov, D., Pearson, A.C., Yen, C.C., Bradley, S.P., and Chudasama, Y. (2025). Cingulate cortex shapes early postnatal development of social vocalizations. eLife 13.

30. Caine, N.G. (1998). Cutting costs in response to predatory threat by Geoffroy’s marmosets (Callithrix geoffroyi). Am. J. Primatol. 46, 187–196.

31. Bishop, M., Weinhold, M., Turk, A.Z., Adeck, A., and SheikhBahaei, S. (2022). An open-source tool for automated analysis of breathing behaviors in common marmosets and rodents. eLife 11.

32. Parshall, M.B., Schwartzstein, R.M., Adams, L., Banzett, R.B., Manning, H.L., Bourbeau, J., Calverley, P.M., Gift, A.G., Harver, A., Lareau, S.C., et al. (2012). An official American Thoracic Society statement: update on the mechanisms, assessment, and management of dyspnea. Am. J. Respir. Crit. Care Med. 185, 435–452.

33. Folschweiller, S., and Sauer, J.-F. (2021). Respiration-Driven Brain Oscillations in Emotional Cognition. Front. Neural Circuits 15, 761812.

34. Kanwal, J.K., Coddington, E., Frazer, R., Limbania, D., Turner, G., Davila, K.J., Givens, M.A., Williams, V., Datta, S.R., and Wasserman, S. (2021). Internal state: dynamic, interconnected communication loops distributed across body, brain, and time. Integr. Comp. Biol. 61, 867–886.

35. Banzett, R.B., Lansing, R.W., and Binks, A.P. (2021). Air hunger: A primal sensation and a primary element of dyspnea. Compr. Physiol. 11, 1449–1483.

36. Ming, X., Patel, R., Kang, V., Chokroverty, S., and Julu, P.O. (2016). Respiratory and autonomic dysfunction in children with autism spectrum disorders. Brain Dev. 38, 225–232.

37. Hardman, C.D., and Ashwell, K.W.S. (2012). Stereotaxic and Chemoarchitectural Atlas of the Brain of the Common Marmoset (Callithrix jacchus) (CRC Press).

